# Molecular Identification of some Indian Muscid flies (Diptera: Muscidae) based on mitochondrial gene COII

**DOI:** 10.1101/208314

**Authors:** Devinder Singh, Ramandeep Achint

**Author notes:** **Corresponding author** - Ramandeep Achint, Department of Zoology and Environmental Sciences, Punjabi University Patiala, Punjab, India (147002).

## Abstract

Taxonomic identification of flies having medical and veterinary importance is often complicated due to the lookalike morphological characters. Molecular identification of five Indian muscid flies (*Musca domestica*, *Musca sorbens*, *Musca crassirostris*, *Stomoxys calcitrans* and *Haematobia irritans*) has been attempted on the basis of mitochondrial COII gene. Sequences of 500-520 bp were analysed and found to be A+T rich. Rate of transitions was higher than transversions. The average haplotype diversity was 0.833 and nucleotide diversity was 0.02547 within the different species, which were calculated with the DnaSP Version 5.0. The genetic distances calculated by K2P model, showed the interspecific distances range more than 8.2%, while the intraspecific distance range had not exceed 0.8%. The 1000 bootstrapped Neighbour-joining tree and Maximum likelihood tree were constructed to establish the phylogenetic relationship between the different muscid species. The results show the robustness of COII gene as a diagnostic marker. The data obtained from this study would be worthy for medical and veterinary entomologists for precise identification of imperative muscid species.

## 1. INTRODUCTION

The flies belonging to the family Muscidae are mostly cosmopolitan in nature. There are approximately 5000 described muscid species under 170 genera and the family is amply represented in all biogeographical regions. They are of great medical and veterinary importance because of their ability to transmit diseases to humans and animals. Adults of many species are passive carrier of pathogens for diseases like typhoid fever, dysentery, anthrax and cholera (Kettle, 1995). Moreover, the fly larvae can be found on dead bodies and are used as a forensic indicator for estimating Post-Mortem Interval (PMI) (Sukontason *et al.*, 2007; Preativatanyou *et al.*, 2010). Adults can be predatory, hematophagous, saprophagous or may feed on different types of plant and animal exudates. Some species have biting mouthparts and others are merely scavengers.

Sometimes the identification of muscid species can be puzzling because of similar morphological markers and it may not be possible to identify the immature stages of many species (Benecke and Wells, 2001). Deoxyribonucleic acid (DNA)-based methods for species identification may solve these problems especially for scientists not formally trained in taxonomy and can be applied on all life stages including incomplete or damaged samples when morphological characteristics were destroyed (Sperling *et al.*, 1994; Wells and Stevens 2008).

Mitochondrial genome had an outsized impact on entomological genetics (Cameron, 2014). As mtDNA is relatively small, abundant and easy to isolate, it has been the favourite target of early genome sequencing projects and the nucleotide sequence of mtDNA from a large number of species has now been determined (http://megasun.bch.umontreal.ca/). Mitochondrial Cytochrome Oxidase I and II genes are suitable as molecular markers due to high degree of variation that has been reported (Wells and Sperling, 1999; Sperliing *et al.*, 1994). The COII gene has been extensively used to address phylogenetic questions in insects at different taxonomic levels: among orders (Liu and Beckenbach, 1992), within an order (Frati *et al.*, 1997), and more successfully within genus or species groups (Beckenbach *et al.*, 1993; Emerson and Wallis, 1995; Mardulyn *et al.*, 1997). The COII gene is more frequently used in evolutionary studies, population genetics and systematics because of the high degree of variation (Junqueira *et al.*, 2002).

The present work represents an attempt to check efficacy of COII gene for the identification of five muscid species from India. The nucleotide sequences generated during this study were submitted to online database. These sequences will be helpful for the identification of medically and veterinary important *Musca domestica*, *Musca sorbens*, *Musca crassirostris*, *Stomoxys calcitrans* and *Haematobia irritans* flies in the future studies.

## 2. MATERIALS AND METHODS

### 2.1 Fly sampling

All individuals were collected with the help of sweeping type net, mainly from four districts of Punjab (Ludhiana, Patiala, Moga and Fatehgarh Sahib), from March to September 2016. Once the flies were caught, they were killed by using ethyl acetate charged killing jar. This was then followed by proper pinning and labelling of specimens. The labels provided information like location, date and timing records as well as other valuable information about the specimens. The flies were identified morphologically using keys given by Emden (1965) and Fonesca (1968). Legs were carefully dissected out from specimens and preserved in absolute alcohol at −20°C for further studies.

### 2.2 DNA isolation, amplification and sequencing

Genomic DNA was extracted using the QIAamp DNA Tissue Kit (QIAGEN) and HiPurA Insect DNA Purification Kit (HiMEDIA). A set protocol was followed with these extraction kits to make sure maximum accuracy of the procedure. Only the insect legs were used for extraction purposes to avoid any possibility of amplifying endosymbiotic microbes associated with the gut of the insect, which could result in wrong interpretation.

Primers forward C2-J-3138 (5′AGAGCTTCTCCTTTAATGGAACA3′) and reverse C2-N-3686 (3′CAATTGGTATAAAACTATGATTTG5`) (Simon *et al*. 1994) were used to amplify the mitochondrial COII gene. The PCR reaction included 0.25μl of each primer, 1.20μl of dNTP, 0.25μl of MgCl_2_, 0.2μl of Phusion DNA polymerase, 1.25μl of stabilizer buffer, 0.8μl of Nuclease-free water and 6μl of DNA adjusted to a final volume of 10μl.

The amplification program had an initial denaturation step at 98°C for two minutes, followed by 38 cycles of 98°C for 30 seconds each, annealing at 49°C for 40 seconds, elongation at 72°C for a minute and a final extension step was added for seven minutes at 72°C. After amplification the PCR products were run on agarose gel for checking up the amplification. After attaining the results from the gel, the amplified samples were sent to SciGenom Labs, Cochin (Kerala) for DNA sequencing. The sequences were then submitted to the GenBank and unique accession numbers were obtained for all these sequences.

### 2.3 Analysis of obtained COII sequences

Once the sequences were obtained, these were analyzed, trimmed and edited using BankIt. This is must to remove unwanted base pairs or stop codons which are included by default, because, these do not allow accurate submissions. Only well-edited sequences were submitted to the GenBank. Table 1 enlists the accession numbers obtained from GenBank for various samples.

The sequences obtained during the present study were compared with already submitted sequences with NCBI database, to calculate intraspecific and interspecific distances. Accession numbers of those downloaded sequences are given in Table 2.

The sequences were aligned by using ClustalW software. Analysis of nucleotide composition, overall transition/transversion ratio (ts/tv), conserved, variable and parsimony informative sites and pairwise nucleotide distances were calculated using MEGA 7. The haplotype diversity (Hd), nucleotide diversity (pi) and Tajima’s *D* values were calculated by DnaSP Version 5.0 (Librado and Rozas, 2009). To reveal the lineage history Maximum Likelihood tree and Neighbour-Joining tree method with 1000 replications in the bootstrap test were used. Codon positions included were 1^st^+ 2^nd^+ 3^rd^+ Non-coding and there were a total of 480 positions in the final dataset.

## 3. RESULTS AND DISCUSSION

### 3.1 Base composition

The conserved, variable and parsimoniously informative sites have been analyzed for the studied species. The overall numbers of conserved sites were 365 and variable sites were only 115. This clearly indicates that COII gene is highly conserved. Out of the 115 variable sites, 96 sites were found to be parsimoniously informative.

In the present study, both nucleotide sequence and the particular nucleotide percentage have been evaluated as these factors are important for studying the variation among different species. The average percentage of each nucleotide for the studied fragment of COII gene was observed to be T= 39.40%, A= 32.95%, C= 13.58% and G= 14.07%. This clearly shows that A+T content of the nucleotide sequences was higher than G+C. Higher A+T frequency is distinguishing make-up of insect mtDNA (Lunt and Hyman, 1997; Bernasconi *et al.*, 2000; Harvey *et al.*, 2003; Junqueira *et al.*, 2004). Table 3 depicts the base composition (in percent) in the different species of Muscidae at the three codon positions. The A and T were dominant at the first position. The G content at first codon position was less than 2% among all the species while in *Stomoxys calcitrans* it was 0%.

### 3.2 Interspecific and intraspecific distance

The difference between intra and interspecific DNA variation has been used to recognize new species for some taxa (Hung *et al.*, 1999). The interspecific distance between Muscid spp. ranged from 8.2% to 13.7%. Pairwise genetic distance (in percent) of muscid species is given in Table 4. The maximum interspecific variation was found in species *Musca crassirostris* and *Stomoxys calcitrans*, then *Musca domestica and Stomoxys calcitrans*, which indicate the efficiency of COII sequence to identify the species. Intraspecific distance was observed between the range 0-0.8% with an average of 0.4% for *Musca crassirostris*, 0% for *Haematobia irritans*, 0.5% for *Stomoxys calcitrans*, 0% for *Musca domestica* and 0.4% for *Musca sorbens*. No variation was detected between the individuals of *Musca domestica* from different regions and *Haematobia irritans* from the same region.

The transition/transversions bias for the present study was *R*=1.108 calculated by Maximum Composite Likelihood Method. The transitions/transversion rate ratios are *k*_*1*_ = 2.166 (purines) and *k*_*2*_ = 3.276 (pyrimidines). The rate of transitions was higher than the rate of transversions. The most frequent transitions were T→C type. The number of transitional substitution was found maximum among *Musca crassirostris* and *Stomoxys calcitrans* and minimum within *Musca domestica* and *Haematobia irritans*. Haplotype diversity (Hd) was within the range from 0.400 to 1.00 (0.400 for *Haematobia irritans*, 0.833 for *Musca crassirostris*, 1.000 for *Stomoxys calcitrans*, 1.000 for *Musca sorbens*, 0.986 for *Musca domestica*) with an average of 0.833. Nucleotide diversity (pi) were calculated with ranges from 0.0041 to 0.5736 (0.00917 for *Haematobia irritans*, 0.04375 for *Musca crassirostris*, 0.01292 for *Stomoxys calcirans*, 0.00417 for *Musca sorbens,* 0.5736 for *Musca domestica*) with average of 0.02547. The Tajima’s *D* values were calculated for studied species, which were all negative. Average frequencies (in percent) of amino acids for the five species of the present study were observed from 0 to 18.3 as shown in figure 2. Maximum frequency 18.3% of Serine and minimum frequency 0% of alanine, asparagines and glutamine were observed.

**Figure 1:**
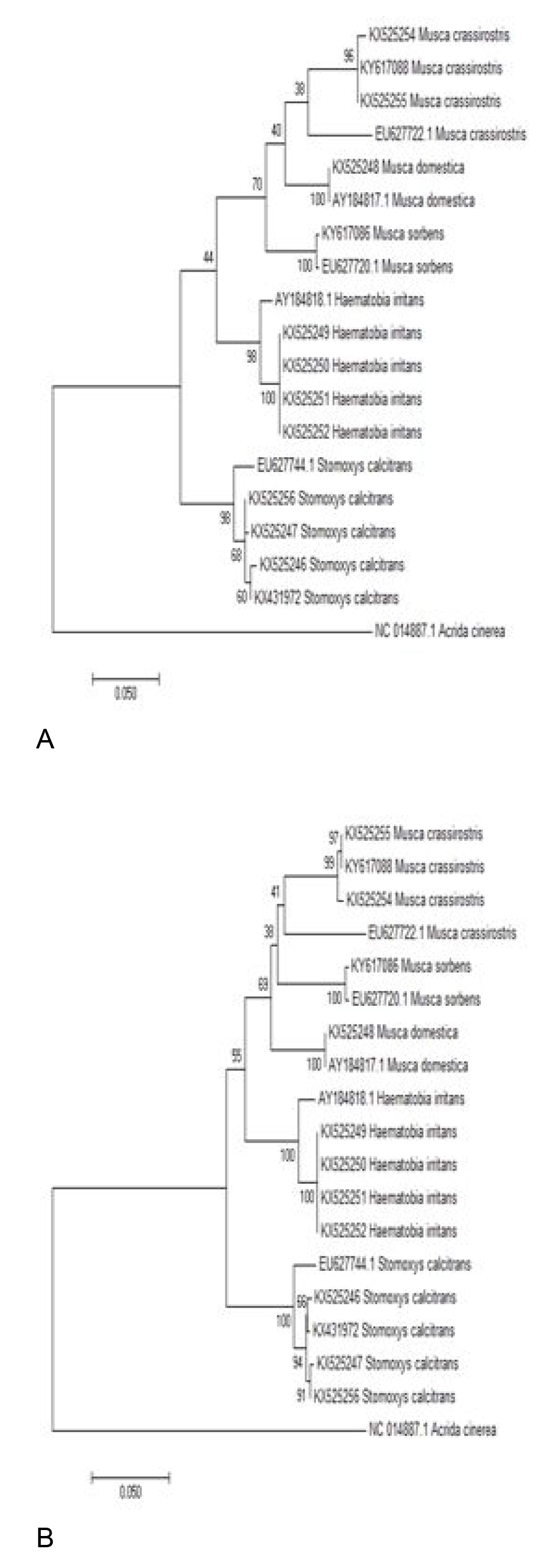
The 1000 bootstrapped A) Maximum likelihood tree and B) Neighbour joining tree under the Kimura 2-parameter model based on COII region sequence of five muscid flies, rooted with *Acrida cinerea*.

**Figure 2:**
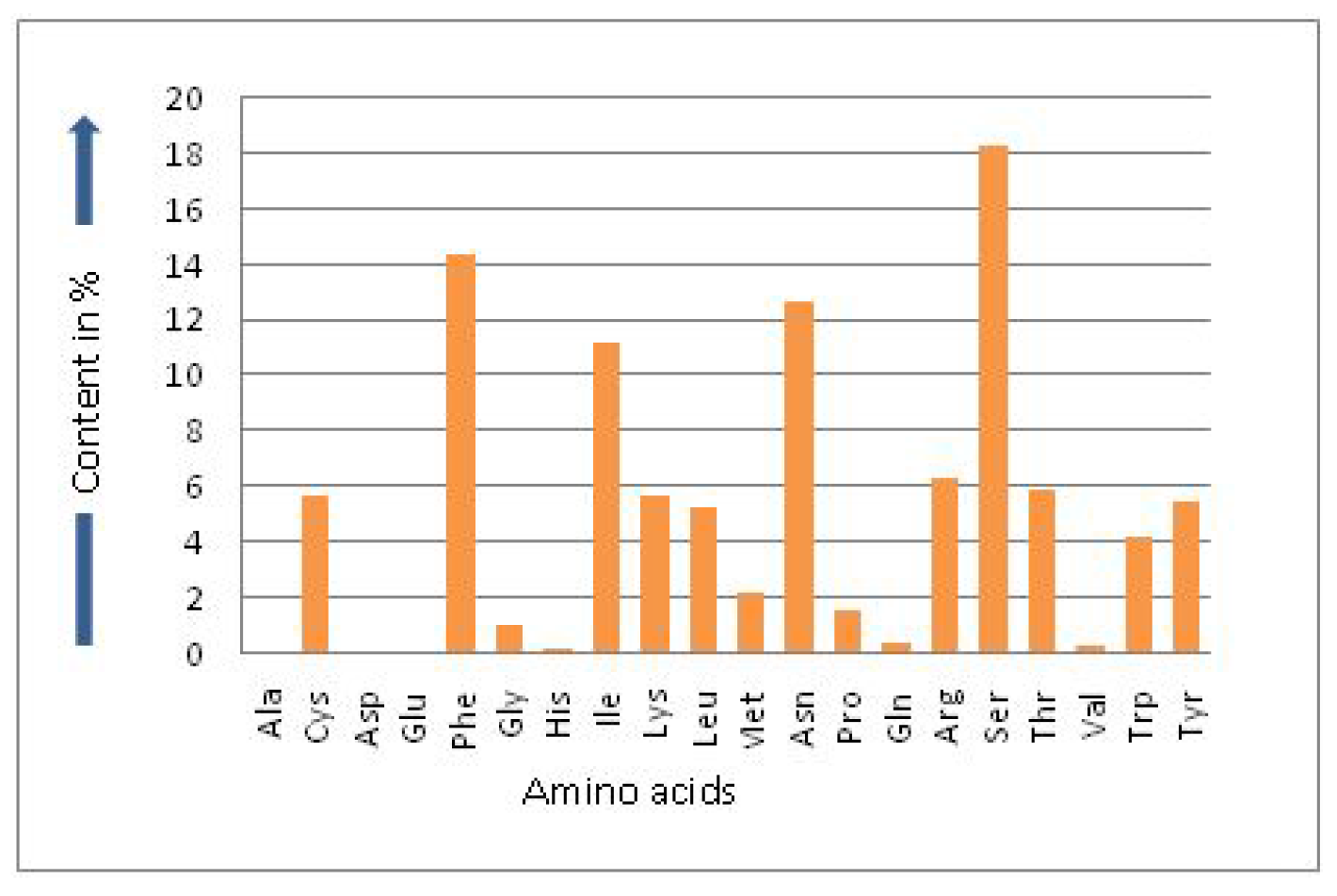
Amino acid content in percent.

### 3.3 Phylogentic Analysis

Sequences of COII region of Muscid flies obtained during this study together with sequences available from NCBI were used for evaluating the evolutionary relationship where *Acrida cinerea* (an orthopteran species) was used as an outgroup for the analysis. Maximum-Likelihood analysis and Neighbour-Joining analysis with 1000 bootstrap by Kimura 2-parameter model showed similar topologies. Nodes linking sequences of individuals of same muscid species had a high bootstrap value. In both analyses, all muscid species separated out correctly and formed a monophyletic clade. The tree clustered into three subclades: first consisting of *Stomoxys calcitrans*, second consisting of *Hamatobia irritans* and third consisting of *Musca sorbens*, *Musca domestica* and *Musca crassirostris*.

## 4. DISCUSSION

It is for the first time that sequence analysis of these 5 muscid species collected from Punjab (India) has been done on the basis of mt COII gene. According to Meyer and Paulay (2005), genetic diversity within the species should not be greater than divergence between the species. For authentic DNA-based species identification, the studied taxa should have less than 1% intraspecific and more than 3% interspecific genetic diversity (Wells and Sperling, 2001). Results presented here reveal a pattern of low intraspecific and high interspecific variations for COII gene in muscid species which totally agreed with the previous studies. Aly *et al*. (2012) studied the interspecific and intraspecific variation of 2 muscid species i.e. *Musca domestica* and *Musca autumnalis*, which was 0% and 12% respectively. Szalanski and Owens (2003) also calculated the interspecific divergence for 5 muscid species in range from 8% to 12%. The average haplotype diversity was 0.833 and nucleotide diversity was 0.025 in the studied species. Low *et al*. (2014) also calculated haplotype diversity between the range 0.733 to 0.961 and nucleotide diversity in range 0.0005 to 0.0014. The negative values of Tajima’s *D* test showed the very low frequency mutations in populations. The genetic distances among the three *Musca* species were also longer than the *Haematobia* and *Stomoxys* species. *Musca domestica* is more closely related to *Musca crassirostris* as compared to *Musca sorbens* as depicted by phylogenetic trees. *Musca* species form a sister clade with *Haematobia irritans*, whereas *Stomoxys calcitrans* form a different clade. Iwasa and Ishiguro (2010) also reported *Haematobia* and *Musca* as sister clade.

The aim of this study was to assess the suitability of mitochondrial COII gene for species level identification of medically and veterinary important muscid flies from India. The current data set provides a useful reference as a backbone for further work, based on COII gene from India. In conclusion, COII gene can be used as a genetic marker to differentiate medically and veterinary important muscid fly species. Due to absence of reference sequences, future studies should be undertaken to generate mitochondrial COII gene reference database for molecular identification of muscid flies from different regions of India.

## References

Aly, S.M., Wen, J., Wang, X. and Cai, J., 2012. Cytochrome oxidase II gene “short fragment” applicability in identification of forensically important insects. Romanian Journal of Legal Medicine, 20: 231–236.

Beckenbach, A. T., Wei, Y. W., and Liu, H., 1993. Relationships in the Drosophila obscura species group inferred from mitochondrial cytochrome oxidase II sequences. Molecular Biology Evolution, 10: 619–634.

Benecke, M. and Wells, J.D., 2001. DNA techniques for forensic entomology. In Forensic Entomology: Utility of Arthropods in Legal Investigations. Byrd. Boca Raton: CRC Press, 341–352.

Bernasconi, M.V., Valsangiacomo, C., Piffaretti, J.C. and Ward, P.I., 2000. Phylogenetic relationships among Muscoidea (Diptera: Calyptratae) based on mitochondrial DNA sequences. Insect Mol. Biol. 9: 67–74.

Cameron, S.L., 2014. Insect Mitochondrial Genomics: Implications for Evolution and Phylogeny. Annual Review of Entomology, 59: 95–117.

Emden, F.I., 1965. The Fauna of India and the adjacent countries, Diptera, Muscidae, 7(I). Govt. Of India, New Delhi, pp. 647.

Emerson, B. C., and Wallis, G. P., 1995. Phylogenetic relationships of the Prodontria (Coleoptera; Scarabaeidae; Subfamily Melolonthinae), derived from sequence variation in the mitochondrial Cytochrome Oxidase II gene. Molecular Phylogenetic Evolution, 4: 433–447.

Fonseca, E.C.M., 1968. Diptera Cyclorrapha Calyptrata: Muscidae. Handbooks for the Identification of British Insects. Royal Entomological Society of London, 10(4b): 118.

Frati, F., Simon, C., Sullivan, J., and Swofford, D. L. (1997). Evolution of the mitochondrial Cytochrome Oxidase II gene in Collembola. Journal of Molecular Evolution, 44: 145–158.

Harvey, M.L. Dadour, I.R. and Gaudieri, S., 2003. Mitochondrial DNA Cytochrome Oxidase I gene: potential for distinction between immature stages of some forensically important fly species (Diptera) in Western Australia. Forensic Science International, 131(3): 134–139.

Hung, S.Y., Chang, C.M. and Yu, T.J., 2006. Determinants of user acceptance of the e Government services: the case of online tax filing and payment system. Government Information Quarterly, 23(1): 97–122.

Iwasa, M. and Ishiguro, N., 2010. Genetic and morphological difference of Haematobia irritans and H. exigua, and molecular phylogeny of Japanese Stomoxyini flies (Diptera:Muscidae). Medical Entomology and Zoology, 61(4): 335–344.

Junqueira, A.C.M., Lessinger, A.C. and Azeredo-Espin, A.M.L., 2002. Methods for the recovery of mitochondrial DNA sequences from museum specimens of myiasis causing flies. Medical and Veterinary Entomology, 16: 39–45.

Juqueira, A.C.M., Lessinger, A.C., Torres, T.T., da Silva, F.R., Vettore, A.L., Arruda, P. and Azeredo-Espin, A.M.L., 2004. The mitochondrial genome of the blowfly Chrysomya chloropyga (Diptera: Calliphoridae). Gene, 339: 7–15.

Kettle, D.S., 1995. Medical and veterinary entomology. Cambridge University Press, pp. 720.

Librado, P. and Rozas, J., 2009. DnaSP v5: A software for comprehensive analysis of DNA polymorphism data. Bioinformatics, 25: 1451–1452.

Liu, H., and Beckenbach, A. T., 1992. Evolution of the mitochondrial Cytochrome Oxidase II gene among 10 Orders of insects. Molecular Phylogenetic Evolution, 1: 41–52.

Low, V.N., Tan, T.K., Lim, P.E., Domingues, L.N., Tay, S.T., Lim, A.L., Goh,T.Z., Panchadcharam, C., Bathmanaban, P. and Sofian-Azirun, M., 2014. Use of COI, CytB and ND5 genes for intra and interspecific differentiation of Haematobia irritans and Haematobia exigua. Veterinary Parasitology, 204(2014): 439–442.

Lunt, D.H. and Hyman, B.C., 1997. Animal mitochondrial DNA recombination. Nature. 387: 247.

Mardulyn, P., Milinkovitch, M. C. and Pasteels, J. M., 1997. Phylogenetic analyses of DNA and allozyme data suggest that Gonioctena leaf beetles (Coleoptera; Chrysomelidae) experienced convergent evolution in their history of host-plant family shifts. Systematic Biology, 46: 722–747.

Meyer, C.P. and Paulay, G., 2005. DNA barcoding: error rates based on comprehensive sampling. Public Library of Science Biology 3, e422 (10 pp.).

Preativatanyou, K., Sirisup, N., Payungporn, S., Poovorawan, Y., Thavara, U., Tawatsin, A., Sungpradit, S. and Siriyasatien, P., 2010. Mitochondrial DNA based identification of some forensically important blowflies in Thailand. Forensic Science International, 202(3): 97–101.

Simon, C., Frati, F., Beckenbach, A., Crespi, B., Liu, H.and Flook, P. 1994. Evolution, weighting and phylogenetic utility of mitochondrial gene sequences and a complilation of conserved polymerase chain reaction primers. Annals of Entomological Society of America, 87:651–701.

Sperling, F.A.H., Anderson, G.S. and Hickey, D.A., 1994. A DNA-based approach to the identification of insect species used for postmortem interval estimation. Journal of Forensic Science, 39(2): 418–427.

Sukontason, K., Sukontason, K.L., Ngern-Klun, R., Sripakdee, D. and Piangjai, S., 2007. Differentiation of the Third Instar of Forensically Important Fly Species in Thailand. Annal of the Entomological Society of America, 97(6): 1069–1075.

Szalanski, A.L. and Owens, C.B., 2003. Sequence Change and Phylogenetic Signal in Muscoid COII DNA Sequences. DNA Sequence, 14(4):331–334.

Wells, J.D., and Sperling, F.A.H., 2001. DNA-based identification of forensically important Chrysomyinae (Diptera: Calliphoridae). Forensic Science International, 120:110–115.

Wells, J.D. and Stevens, J.R., 2008. Application of DNA based methods in forensic entomology. Annual Review of Entomology, 53: 103–20.

Wells, J.D. and Sperling, F.A., 1999. Molecular phylogeny of Chrysomya albiceps and C. rufifacies (Diptera: Calliphoridae). Journal of Medical Entomology, 36(3): 222–226.

